# Recombinant production of a functional SARS-CoV-2 spike receptor binding domain in the green algae *Chlamydomonas reinhardtii*

**DOI:** 10.1101/2021.01.29.428890

**Authors:** A. Berndt, T. Smalley, B. Ren, A. Badary, A. Sproles, F. Fields, Y. Torres-Tiji, V. Heredia, S. Mayfield

**Affiliations:** Division of Biological Sciences, University of California, San Diego, La Jolla, California, USA

**Author notes:** Corresponding author *E-mail address:* *Mailing address:* University of California, San Diego 9500 Gilman Drive MC#0116 La Jolla, CA 92093.

## Abstract

Recombinant production of viral proteins can be used to produce vaccine antigens or reagents to identify antibodies in patient serum. Minimally, these proteins must be correctly folded and have appropriate post-translation modifications. Here we report the production of the SARS-CoV-2 spike protein Receptor Binding Domain (RBD) in the green algae *Chlamydomonas.* RBD fused to a fluorescent reporter protein accumulates as an intact protein when targeted for ER-Golgi retention or secreted from the cell, while a chloroplast localized version is truncated, lacking the amino terminus. The ER-retained RBD fusion protein was able to bind the human ACE2 receptor, the host target of SARS-CoV-2, and was specifically out-competed by mammalian cell-produced recombinant RBD, suggesting that the algae produced proteins are sufficiently post-translationally modified to act as authentic SARS-CoV-2 antigens. Because algae can be grown at large scale very inexpensively, this recombinant protein may be a low cost alternative to other expression platforms.

## INTRODUCTION

In late 2019 a novel coronavirus was identified in the Wuhan province of China with a genome sequence closely resembling that of the Severe Acute Respiratory Syndrome (SARS) coronavirus identified in 2003. Thus the novel coronavirus was named SARS-CoV-2. The respiratory disease the virus causes has since been named COVID-19 (COronaVIrus Disease 2019). The virus quickly spread worldwide and was classified as a global pandemic in April of 2020 (Cucinotta and Vanelli, 2020), and has continued to spread almost 1 year after its initial identification.

Widespread use of nucleic acid tests that detect the SARS-CoV-2 RNA genome, such as RT-qPCR have become the standard method to detect viral infection. However, laboratory assays that measure antibody responses and determine seroconversion are not yet comparatively as widely available. While such serological assays are not well suited to detect acute infections, and indeed antibody production lags days behind symptoms and infectiousness, multiple relevant applications exist for such antibody tests as they are one of the best indicators of prior immune responses to viral antigens, and therefore might indicate individuals are immune (Carter et al., 2020). In addition to use as a reagent to identify serum antibodies, recombinant spike protein can also function as a vaccine antigen, and nearly all of the SARS-CoV-2 vaccines being deployed today are based on the SARS-CoV-2 spike proteins. As these vaccines are deployed at a population scale, monitoring short- and long-term immune response to the vaccine target (the SARS-CoV-2 spike (S) protein) will be a key component of characterizing vaccine efficacy, but will also potentially provide information of when booster-doses of the vaccines might be required, as immune responses can wane over time, while the virus is still known to be circulating in the population. This may become especially important as practical adherence to vaccination schedules established during clinical trials and “mix and match” use of different vaccines becomes a practical reality. To develop such antibody tests, it is critical to produce the viral protein antigens at extremely large scale and at an affordable price, so that antibody tests can become available around the world, and not just in the economically advantaged countries.

The spike protein of SARS-CoV-2 mediates viral entry into host cells by first binding to the host angiotensin-converting enzyme 2 (ACE2) receptor followed by fusion of the viral and host membranes and release of the viral RNA into the host cell (Lan et al., 2020; Li et al., 2005). The receptor-binding domain (RBD) of the spike protein is located in the S1 subunit and directly mediates the interaction with the ACE2 cell receptors (Lan et al., 2020). The SARS-CoV-2 RBD is the protein primarily used in COVID-19 vaccines, and patients that have survived the infection often have serum antibodies directed against this viral protein domain (Tai et al., 2020). Antibody neutralization assays have also identified a strong correlation between high levels of serum RBD specific binding antibodies and viral neutralization (Premkumar et al., 2020). Based on this collective data, the SARS-CoV-2 RBD has become the antigen most used for serological assays in addition to the antigenic target of multiple vaccines.

Production of useful SARS virus spike proteins and their subdomains has been a challenge for non-animal cell production systems. Production of the 2003 SARS virus RBD or full spike protein utilizing an *E. coli-based* system, showed poor protein folding and aggregation into inclusion bodies. The resulting protein was much less immunogenic than the same protein produced in mammalian or insect cells (Chen et al., 2005; Du et al., 2009). Production of corona virus RBD proteins in fungal systems such as *Pichia pastoris* have demonstrated potential, but the productivity tends to be orders of magnitude lower than what these yeast systems have been shown to be capable of for other recombinant proteins (Chen et al., 2017, 2014).

Previous studies have shown success in plant-based systems for recombinant production of viral recombinant proteins (Demurtas et al., 2016; He et al., 2014; Meyers et al., 2008). The membrane (M) and nucleocapsid (N) proteins of SARS were expressed transiently in *Nicotiana benthamiana*, with the N protein demonstrating antibody recognition in convalescent serum samples (Demurtas et al., 2016). The main drawback of plant-based expression systems is that they have low biomass productivity compared to microbial systems, and have laborious and technically demanding transformation methods that can require months to achieve (Muthamilselvan et al., 2019).

Green microalgae can be grown photosynthetically or heterotrophically and can scale very rapidly (Fields et al., 2018). Algae have been demonstrated to fold complex eukaryotic proteins (Rasala et al., 2012, 2010), to be amenable to sophisticated molecular genetic tools (Sproles et al., 2021), and to express recombinant proteins which can be directed to any subcellular structures (Rasala et al., 2013). Collectively this allows for the rapid production of complex proteins that can be grown at large scale in a cost effective manner (Rosales-Mendoza et al., 2020).

Here we examined the potential of utilizing the unicellular green microalgae *Chlamydomonas reinhardtii* as a production platform for recombinant SARS-CoV-2 spike RBD protein. We tested three protein targeting strategies for recombinant protein production within *C. reinhardtii*, by appending different intracellular localization motifs to the transgene. The RBD proteins were targeted either to be retained in the Endoplasmic Reticulum-Golgi pathway, secreted out of the cell into the culture media, or targeted for accumulation within the chloroplast. We found that recombinant RBD targeted to the chloroplast accumulated to high levels, but appeared to be truncated by ~6 kD at the amine end of the mature protein, and this protein was not recognized by anti-RBD antibodies in western blotting assays. RBD proteins targeted to the ER or secreted from the cell produced a protein of the expected size and correct amino acid sequence. We purified spike RBD protein from the ER-Golgi retained version and demonstrated that it specifically binds to recombinant ACE2 protein at a similar affinity as mammalian expressed spike-RBD. These data demonstrate the potential of using eukaryotic algae as an efficient and scalable platform to make correctly folded and functional spike-RBD recombinant proteins that could be used in large scale antibody assays or as potential vaccine antigens.

## RESULTS

### Design of SARS-Covid2 spike protein RBD expression cassette for recombinant protein production in algae

Based on previous studies, as well as bioinformatics and structural modeling, we elected to produce amino acids 319-537 of the SARS-CoV-2 spike protein comprising the RBD (Wrapp et al., 2020). For high throughput screening to rapidly identify recombinant protein production, we fused this domain to the N-terminus of the green fluorescent protein derivative mClover (Lam et al., 2012). The *C. reinhardtii* nuclear genome codon optimized fusion genes of the spike-RBD and mClover were then placed in a modified expression vector based on components of the previously published pBR9 and pOpt vectors (Rasala et al., 2013, Lauersen et al., 2015) (Figure 1 A). Expression was driven with the semi-synthetic AR1 promoter (Rasala et al., 2012). A 5’ Bleomycin resistance gene *(ble)* was included as part of the expression cistron to allow for selection of high expressing clones on Zeocin containing media, while a foot and mouth disease 2A ribosomal-skip motif was placed between the *ble* coding region and the RBD::mClover fusion protein, so that the RBD::mClover accumulated as a single fusion protein (Rasala et al., 2012). A separate Hygromycin resistance gene driven by a beta-tubulin promoter was placed 3’ of the RBD transgene to allow for double selection, ensuring that the RB::mClover transgene is intact in any transformant selected (Berthold et al., 2002).

**Figure 1.**
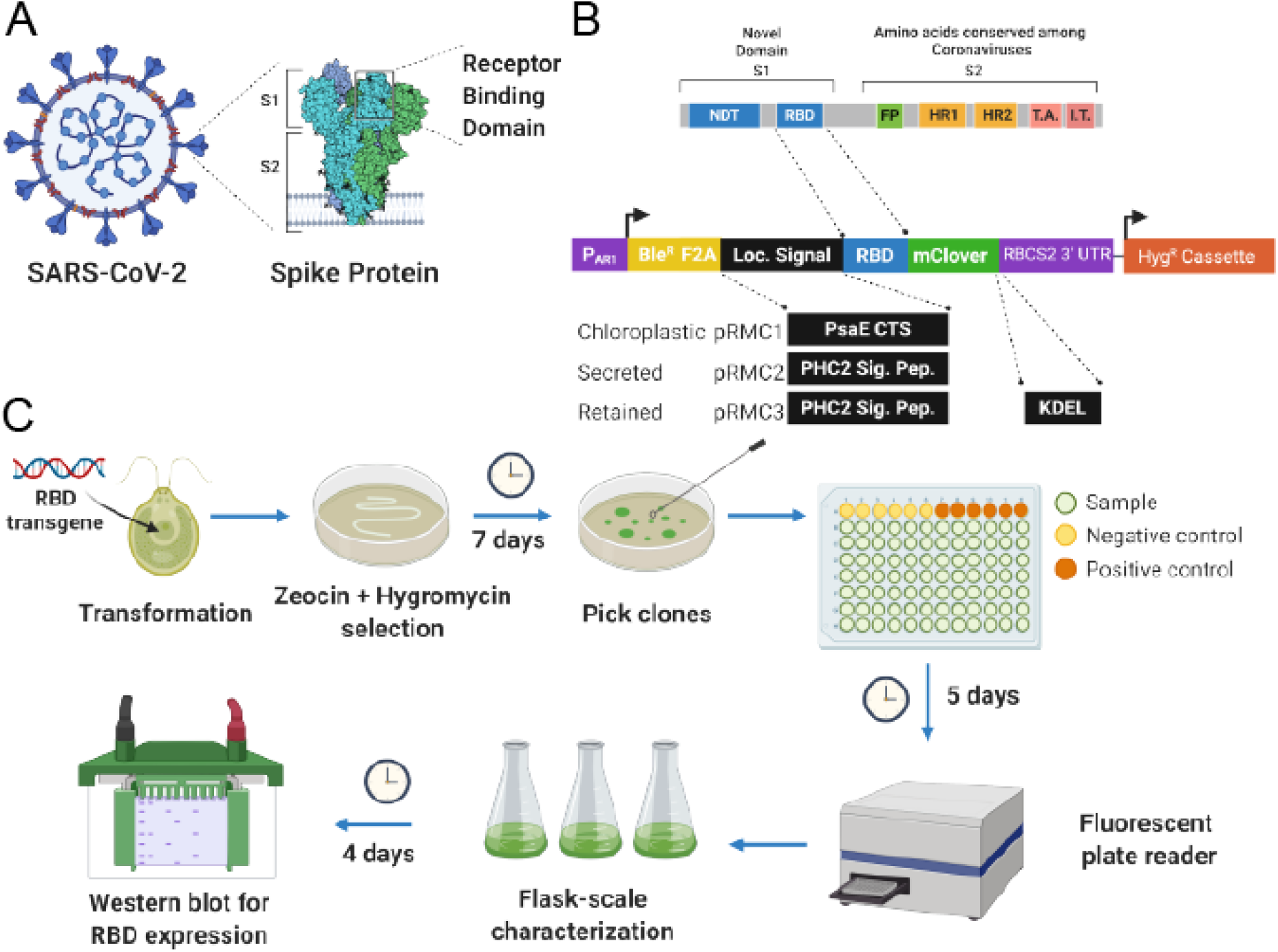
(A) Diagram of the SARS-CoV-2 viral particle with crystal structure of the spike (S) protein highlighted with Subunits 1 and 2 indicated (S1, S2, respectively). The Receptor Binding Domain (RBD) is located in the more variable subunit 1. (B) Vector design and construction. The peptide structure of the spike protein indicating the N-Terminal Domain (NTD), Receptor Binding Domain (RBD), Fusion Peptide (FP, Homology region 1 and 2 (HR1, HR2), Transmembrane association domain (TA) and Intracellular Terminal (IT). A *C. reinhardtii* nuclear codon optimized version of the RBD-coding sequence was cloned in to a vector containing the AR1 promoter (P_AR1_) driving a transcriptional fusion of the Bleomycin resistance gene (BleR), FMDV Foot-and-mouth disease virus 2A (F2A) ribosomal-skip motif and 5’ mClover green fluorescent protein tag. A separate Beta-tubulin2 promoter driving Hygromycin resistance was used for secondary selection. Three different versions of the RBD were generated. A chloroplast-directed version through N-terminal fusion of the PsaE chloroplast transit sequence, a secreted version by the addition of the PHC2 secretion signal peptide, and an ER-Golgi system retained version by the subsequent addition of a C-terminal KDEL Golgi retention sequence. (C) Schematic summarizing transformation process and timeline including drug selection, clone down selection through 96-well microtiter plates, and then flask-scale characterization of candidate RBD-expressing lines.

### Targeting of the RBD::mClover protein to subcellular compartments in transgenic algae

Since it is known that there are different protein folding and posttranslational modifications made to proteins depending on which organelle a protein is directed to, we generated three versions of the RBD::mClover construct; 1) A chloroplast directed version, produced by adding the psaE chloroplast transit sequence to the N-terminus of the RBD fusion protein, 2) a secreted version, produced by adding Pherophorin 2 (PHC2) signal peptide to the N-terminus of the RBD fusion protein, and 3) a ER-Gogli retained version produced by the addition of a C-terminal KDEL retention motif to the carboxy end of the RBD fusion protein containing the PHC2 secretion peptide (Figure 1 B).

### Transformation and high throughput screening for recombinant protein production

The three vectors were linearized and transformed separately into algae via electroporation. Following recovery on complete media, cells were selected on media containing both Zeocin and Hygromycin. Ten days post-transformation, individual colonies were picked into 96-well microtiter plates containing TAP media and grown for two days. The clones were then passaged at a 1:4 dilution in to fresh TAP media for two days and fluorescence analysis, using a plate reader, was used to identify strains with high mClover expression. The mClover fluorescence signal was normalized to chlorophyll fluorescence and compared to both the CC124 starting strain and a previously characterized GFP-expressing strain (Fields, 2019). Several hundred mClover expressing transformants were recovered for both the secreted and ER retained clones, from three independent transformations, while only a few dozen colonies were recovered from the chloroplast targeted strains, despite the same amount of DNA being used in each of the transformations. Similarly, by mClover fluorescence analysis, 10-30% of all secreted and ER retained transformants showed fluorescence well above wild type, while only about 1-5% of the chloroplast-directed strains showed any mClover fluorescence.

### Characterization of RBD::mClover localized to different compartments

When characterized by SDS-PAGE followed by Western Blotting using anti-GFP antibodies to detect the mClover tag, we found cell pellets of the secreted and ER retained versions of the RBD::mClover protein generated a band at the expected molecular weight of ~51 kDa (Figure 2). Similarly, ammonium sulfate precipitated protein from the media of the RBD::mClover secreted construct had a detectable band at ~51 kDa. Of the few recoverable Chloroplast-directed RBD::mClover transformants, all showed a smaller anti-GFP immunoreactive product at about 45 kDa, clearly smaller than the expected 51kDa product. Further characterization using anti-SARS-CoV-2 RBD polyclonal antibodies to probe the western blots, revealed that while the secreted and ER-Golgi retained proteins were detected as expected 51 kDa bands, the chloroplast directed proteins at ~45 kDa were not detected, despite generating comparatively stronger anti-GFP signal (Figure 2). It should be noted that a cross-reactive band at ~51kDa is observed in the wild-type algae lysate when blots are probed with the anti-RBD Rabbit Polyclonal antibody from Sino Biological. When equal amounts of protein were loaded, darker bands were observed in lysates from the ER-Golgi retained and secreted RBD::mClover strains at the expected molecular weight compared to either the wild type or Chloroplast directed strain (See Supplementary Figure 1 for loading control verification using anti-AtpB).

**Figure 2.**
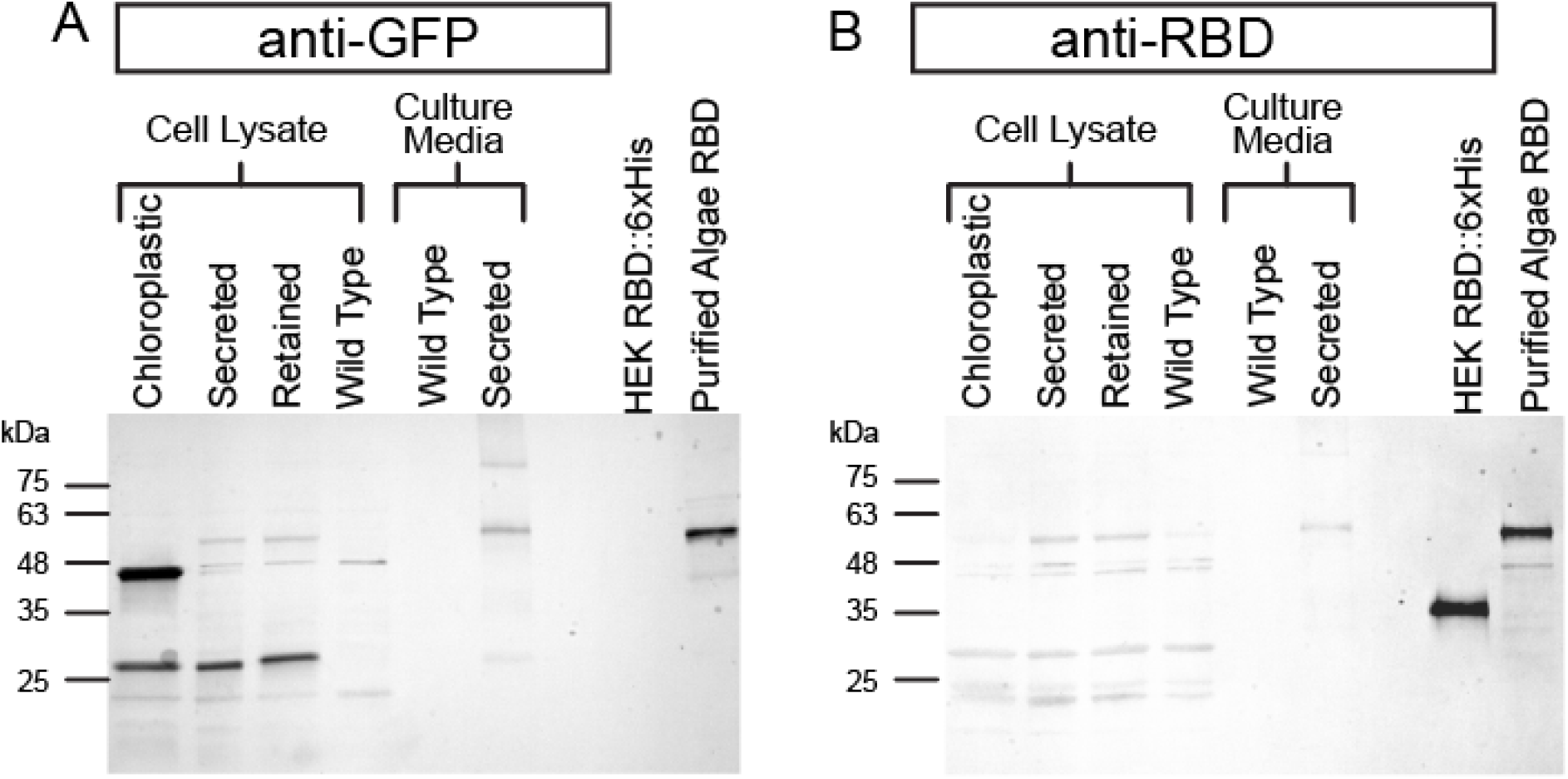
(A) Western blot characterizing the expression of RBD::mClover in the cell pellet lysate or secreted supernatant of transformants from vectors pRMC1/2/3 by probing the blot with anti-GFP. 5 ug of total protein from the cell pellets were loaded in to each well while a 10 uL of 1:200 concentration of the cell culture supernatant or 5 uL of partially purified Algae RBD::mClover was loaded. 100 ng of HEK-cell produced RBD::6xHis tag was used as a control. A background band is observed at ~45 kDa in all samples including our wild-type control. The expected size of all RBD::mClover gene products is 51kDa. The Chloroplast-directed version of our RBD construct the main product appears as a strong band below the 48kDa molecular weight marker while the Secreted or ER-Golgi retained version appears above the 48kDa marker. (B) A western blot with the same organization as before but probed with Rabbit anti-RBD polyclonal antibody. The Chloroplast-directed version of RBD::mClover does not appear to be detected while the Secreted and ER-Golgi retained versions are. A background band is observed in all samples including the wild type control at ~51 kDa.

**Figure 3.**
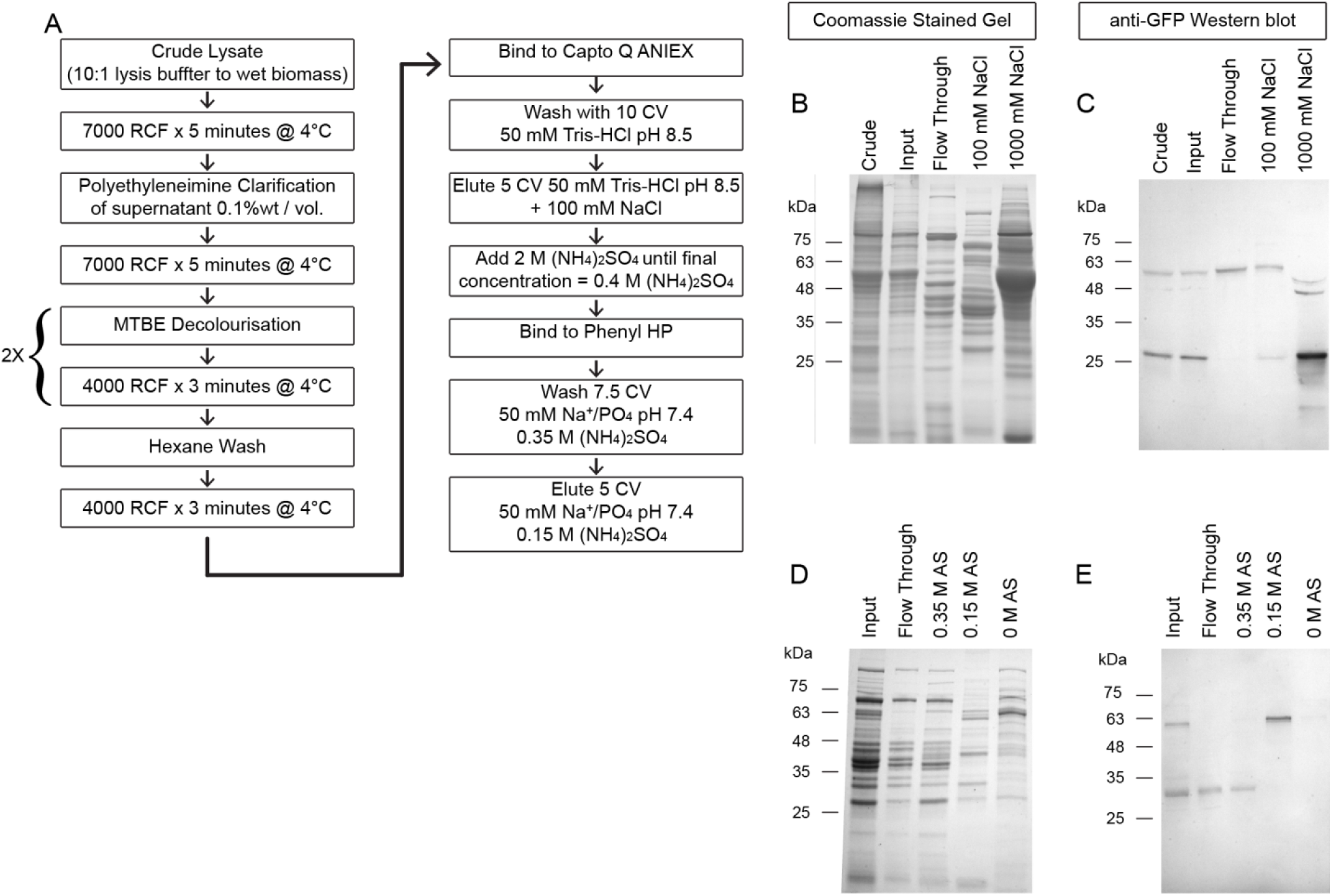
(A) Schematic summarizing our clarification and decolorization protocol prior to column chromatography purification of the RBD::mClover domain through successive Anion Exchange and Hydrophobic interaction resin chemistries. (B) Coomassie stained gel of crude lysate total lysate, Polyethyleneimine/Organic solvent clarified input fraction followed by the Capto Q chromatography Flow through, 100 mM NaCl elution of the RBD::mClover, and 1 M NaCl stripping of the column. Crude and input represent 10 uL of lysate while other fractions represent the same volume of a 20X concentration from the chromatography eluted fractions. (C) Western blot characterization of the same samples as in panel B but using anti-GFP antibody to probe for the RBD::mClover Complex. (D) Coomassie stained gel characterizing the Hydrophobic-interaction purification of the anion-exchange 100 mM NaCl fraction. All samples represent 10 uL of a 20X concentration each fraction. The input represents the 100 mM NaCl anion exchange fraction after the addition of Ammonium Sulfate (AS) to 0.4M. (E) Western blot Characterization of the same samples using anti-GFP to detect the RBD::mClover gene product. The RBD::mClover can be separated from the lower molecular weight mClover degradation product.

The chloroplast localized RBD::mClover protein was then purified using anion exchange chromatography followed by anti-GFP magnetic bead immunoprecipitation, and the partially purified protein products were characterized using protein mass spectrometry analysis. The most N-terminal peptide fragment detectable by mass spectroscopy in the chloroplast-directed RBD::mClover corresponded to a protein product that would be 9 kDa smaller than the predicted 51 kDa full length mature protein, while a parallel experiment using the ER-Golgi retained protein identified peptide matches across the entire length of the protein (See Supplemental Figure 2).

### Purification and characterization of the RBD::mClover fusion proteins

To characterize the RBD protein produced using *C. reinhardtii*, we opted to purify the RBD::mClover from the ER-retained version rather the secreted version, because we noticed that the protein had similar molecular mass, suggesting similar post-translational modifications, and because we noticed high molecular weight aggregates in the precipitated secreted proteins. Using a variety of chromatography resins, we were able to purify sufficient amount of ER-retained RBD::mClover to characterize the protein for receptor-interaction activity, and to identify that total RBD::mClover protein was 0.1% of total soluble protein in total cell lysate and 2.6% of total protein after this partial purification. We determined initial and partially purified final recombinant protein yields of 31 μg/gram of wet biomass and 1.8 μg/gram of wet biomass, respectively.

### Functional characterization of algae-produced SARS-CoV-2 spike RBD protein

The SARS-CoV-2 spike RBD interacts with human host cells through ACE2 receptor binding (Lan et al., 2020, p. 2; Tai et al., 2020; Wrapp et al., 2020). To determine if the algae-expressed ER-retained RBD::mClover was correctly folded and functional, we established a human ACE2 receptor binding assay for SARS-CoV-2 RBD-containing proteins. HEK cell line expressed biotinylated soluble human ACE2 receptor was immobilized on Strepavidin coated microtiter plates and used as a substrate for SARS-CoV-2 RBD binding. To validate the assay we first determined that that HEK cell expressed SARS-CoV-2 RBD C-terminally fused to rabbit IgG Fc (RBD::rFc) could bind to the immobilized ACE2, and could identify that the recombinant RBD bound the ACE2 protein at the EC_50_ of ~0.6ng/mL (Figure 4A). This affinity is within the nM range previously reported (Liu et al., 2020) but 10-fold higher than the EC_50_ reported by the manufacturer (2-6 ng/mL). Next we added a constant amount of our partially purified algae-expressed RBD::mClover, and then titrated in the RBD::rFc as a competitor. Using anti-GFP antibodies to detect RBD::mClover bound to the immobilized ACE2 receptor, we showed that the RBD::rFc will compete RBD::mClover binding at an IC_50_ of ~1 μg/mL which is approximately equal to the 2 μg/mL of the algae produced RBD::mClover loaded in to each well (Figure 4). This suggests that the algae-produced SARS-CoV-2 RBD is functional in ACE2 binding, and appears to have a similar affinity as SARS-CoV-2 RBD produced in HEK cells, the standard for functional SARS-CoV-2 RBD protein.

**Figure 4.**
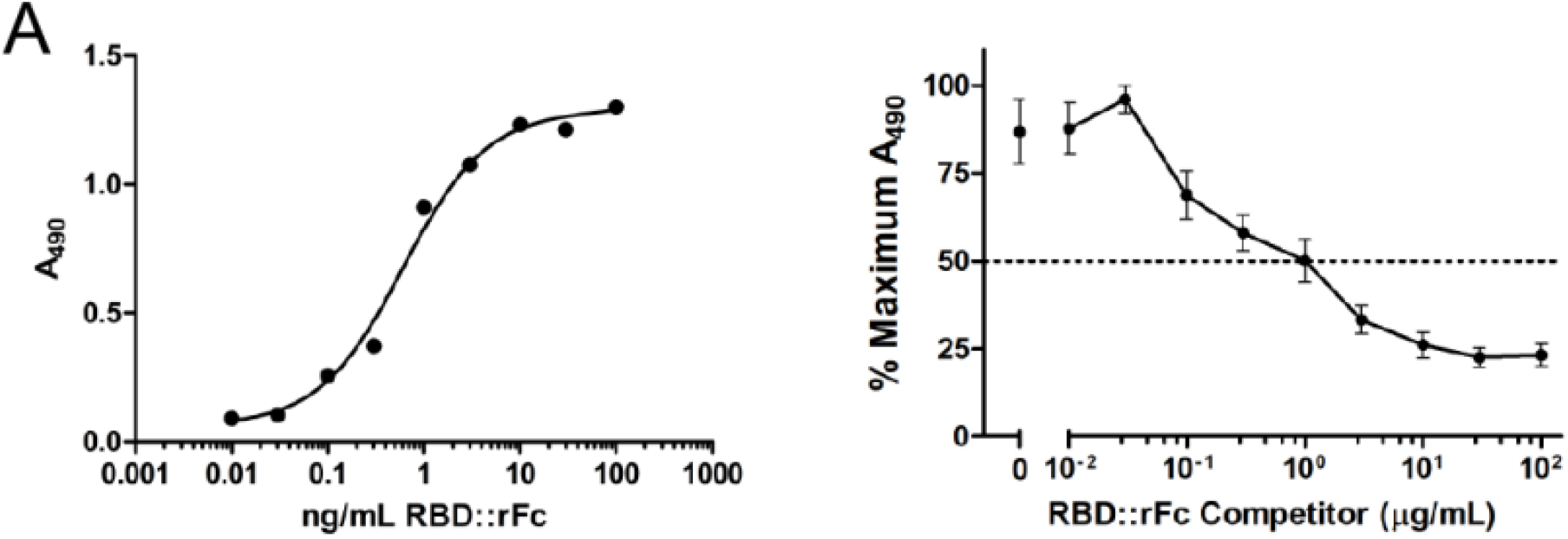
(A) ACE2 Receptor binding assay of HEK-cell produced RBD fragment fused to Rabbit IgG Fc fragment (RBD::rFc). Binding was detected by anti-Rabbit IgG-HRP antibodies. (B) ACE2 Receptor competition assay between a constant concentration of partially purified ER-Golgi Retained Algae-Produced RBD::mClover (2 μg/mL) and increasing amounts of RBD::rFc showing specific competition. RBD::mClover binding was detected using anti-GFP HRP antibodies. Data are shown as mean and standard deviation of normalized A_490_ signal over three independent experiments across three different days.

## DISCUSSION

Here we have demonstrated that production of a correctly folded and functional SARS-CoV-2 spike protein Receptor Binding Domain is possible in the green alga *C. reinhardtii.* By fusing the viral protein to a fluorescent mClover protein, we could use a high throughput fluorescent screening strategy to rapidly identify strains of algae expressing sufficient quantities of SARS-RBD to test protein accumulation and function. We showed that nuclear encoded transgenes, directed to either the ER or secreted from the cell, produce a fusion protein of the expected size that had the correct amino acid sequence, and that appeared to be correctly folded and post-translationally modified, allowing the protein to function in ACE2 receptor binding. Protein targeted to the chloroplast appeared to be truncated, likely by a chloroplast specific protease, resulting in a protein that was not recognized by RBD polyclonal antibodies. Mass spectral analysis suggests that the chloroplast truncated protein is the result of an N-terminal truncation in the chloroplast-directed protein. Using partially purified ER localized algae SARS-CoV-2 RBD to conduct ACE2 receptor binding interaction assays, we could demonstrated that the algae-produced ER-retained version of RBD::mClover did indeed interact in a specific manner with its cognate human host receptor, at a perceived affinity similar to mammalian expressed SARS-CoV-2 RBD, suggesting that the folding and post translational modifications of the recombinant algae RBD are sufficient to mediate appropriate RBD binding activity. Collectively these data demonstrate the suitability of algae as a platform for the rapid production of recombinant proteins that are correctly folded and post-translationally modified to create a functional protein. Because algae can be grown at scale for a fraction of the cost of mammalian cell lines, this system offers the potential to produce this recombinant viral protein, for a variety of uses including as an antigen to detect serum antibodies against SARS-CoV-2 RBD or even as a potential viral antigen for vaccine development, in a rapid and cost-effective manner.

## MATERIALS AND METHODS

### Strain and Culture conditions

*Chlamydomonas reinhardtii* strain CC124 was used throughout experiments. Cells were grown on standard TAP media (Gorman and Levine, 1965) under 24-hr light conditions at ~22-25 °C at a photon flux of 125 μE/m^2^/sec on shaker tables rotating at 110 RPM.

### Vector design & electroporation

#### SARS-CoV2 RBD Sequence design and optimization

The genomic sequence of SARS-CoV-2 (Wuhan-Hu-1 strain) was retrieved from the NCBI (NC_045512). Amino acids 319 to 542 of the spike protein comprising the RBD were codon optimized for *C. reinhardtii* nuclear genes (Amanat et al., 2020). Briefly, we generated a codon-usage table using the mRNA sequences derived from the top 1600 (~10%) nuclear genes expressed in *C. reinhardtii* in TAP media under light growth conditions (Ngan et al., 2015). Codon optimization was then performed using Unipro UGENE (Okonechnikov et al., 2012) with final codon rotations to avoid direct repeats or regions of >70% GC content that were flagged as difficult to synthesize by the IDT gBlock design tool. The full codon optimized sequence was synthetized as a double-stranded gBlock oligonucleotide by IDT (Integrated DNA Technologies Coralville, IA).

#### Expression Vector design and assembly

The expression vector was derived from previously published pBR9 (Rasala et al., 2013), pOPT mClover (Lauersen et al., 2015) and pHyg3 (Berthold et al., 2002). Standard PCR-based amplification to add overlap-adaptors and Gibson-style assembly methods (HiFi Assembly Kit, New England Biolabs, Ipswich, Massachusetts) were used to assemble the final vectors. The entirety of the *C. reinhardtii-specific* payload was sequenced by Sanger Method at Eton Bioscience (San Diego, CA).

Three different sub-cellular localization strategies were chosen for RBD expression. First, a chloroplast-localized version was produced by N-terminal fusion of a previously characterized psaE chloroplast transit sequence localization signal (Franzén et al., 1990) to the RBD domain generating vector pRMC1. The genomic sequence spanning the psaE start codon until four amino-acids after the CTS cleavage site were amplified from *C. reinhardtii* strain CC125 genomic DNA purified by standard Phenol/Chloroform/SDS lysis and Isopropanol/Ethanol precipitation method. The second construct contained a secretion peptide motif (as predicted by SignalP5.0) from the Pherophorin 2 gene (PHC2, Almagro Armenteros et al., 2019). The signal peptide was specifically chosen because PHC2 is known to be abundant in the *C. reinhardtii* secretome (Luxmi et al., 2018). The genomic locus was amplified and placed upstream of the RBD coding sequence, as indicated above for the psaE CTS, to generate vector pRMC2. Finally, an ER-Gogli retained version of the RBD was generated by adding an additional KDEL retention motif to the C-terminus of the RBD coding sequence in pRMC2 to generate pRMC3. All vector maps are included as genbank files in supplementary information.

#### Algae Transformation

All plasmid vectors were prepared by Qiagen midi prep and 10-20 ug of DNA linearized by digest with KpnI-HF and XbaI restriction enzymes at 37°C in NEB Cutsmart buffer with enough units of activity to generate ~5X overdigest in two hours (New England Biolabs, Ipswich, MA). Reaction was stopped and DNA partially purified by NaCl/Isopropanol precipitation followed by three 70% Ethanol washes and resuspension in 1 mM Tris-HCl, 0.1 mM EDTA overnight at 4°C. A starter culture of *C. reinhardti* strain CC124 was grown in standard TAP media to near saturation (~3×10^6^ cells/mL) then diluted back in to 0.25×10^6^ cells/mL in 300 mL of TAP media in a baffled flask ~24 hours before transformation. The next day, cells were collected at a density of 0.75-1.0×10^6^ cells/mL by centrifugation at 3000 rcf x 10 minutes at 16°C in sterile disposable conical bottom centrifuge bottles. The cells were then resuspended in GeneArt^®^ MAX Efficiency^®^ Transformation Reagent for Algae (Thermo Scientific, Waltham, MA) and processed according to manufacturer’s instructions. Each construct was electroporated using a Gene Pulser Xcell Electroporation System (Bio-Rad Laboratories, Hercules, CA) with 2 ug of linearized vector DNA. After over-night recovery in 40 mM Sucrose in TAP media, cells were pelleted, resuspended in 5 mL of fresh TAP media and 200 uL of cells spread on 10 cm diameter plates containing 15 μg/mL Zeocin and 30 μg/mL Hygromycin B (Thermo Scientific, Waltham, MA) in TAP+15 g/L agar media.

#### Transformant Down-Selection

Ten days after electroporation, individual colonies were transferred into 96-well microtiter plates containing liquid TAP media; CC124 strain and previously generated GFP-expressing strain (Fields et al., 2019) were included in each plate as internal positive and negative controls. mClover fluorescence, chlorophyll fluorescence, and optical density were quantified following previously published methods (Fields et al., 2019); mClover fluorescence was normalized to chlorophyll to account for differences in cell density. The seven highest normalized mClover fluorescence clones were then inoculated into 50 mL shaker cultures, and grown for three days until late log phase was achieved (1-3×10^6^ cells/mL), at which point one mL of culture was harvested for further analysis. For pRMC1 and pRMC3 the supernatant was aspirated off and the pellet was snap frozen at −70°C. For pRMC2 the 50 mL of supernatant was recovered, centrifuged a second time to remove any residual cell debris, and then 40 mL of culture media snap frozen. Subsequently, secreted RBD::mClover was precipitated by addition of solid ammonium sulfate to a final concentration of 302 g/L (~50% saturation at 0°C), followed by incubation over night at 4°C with mixing, then centrifugation at 16×10^3^ rcf. The supernatant was removed and the pellet was resuspended in 200 μL of 50 mM Tris-HCl pH 8.5. In preparation for Western Blot analysis, cell pellets were quick thawed in a room temperature water bath, lysed in 100 μL of Bugbuster (Sigma Aldrich, St. Louis, Missouri) for 10 minutes on ice with the addition of Basemuncher (Abcam, Cambridge, MA), cell debris was pelleted (5 minutes × 16×10^3^ rcf at 4°C) and 60 μL of clarified supernatant recovered. Proteins were denatured by addition of 20 μL 4X Laemmli buffer with 10% BME and then heat treated at 80°C for 10 minutes before 20 μL were loaded on to a Tris-Glycine TGX 12% acrylamide gel for SDS-PAGE (Bio-Rad Laboratories, Hercules, California).

#### Western Blotting and Coomassie staining

After separation on acrylamide gels, proteins were transferred to nitrocellulose membranes via semi-dry transfer (15 V x 60 minutes). Membranes were washed 3×2 minutes with TBSMT (50 mM TrisHCl pH 7.4, 150 mM NaCl, 0.05% Tween-20) then blocked for 20 minutes in Haycock’s blocking solution; 1% wt/vol Polyvinylpyrrolidone in TBSMT (Haycock, 1993). Recombinant gene product was either detected with Goat anti-GFP conjugated to Alkaline Phosphatase (ab6661 Abcam, Cambridge, Massachusetts) at 1:5000 dilution or 1:5000 Rabbit Polyclonal anti-SARS-CoV2 RBD (Cat# 40592-T62 Sino Biological, Chesterbrook, PA), in Haycock blocking solution. The latter was then detected with Goat anti-Rabbit::AP at 1:10,000 dilution. All antibodies were incubated for 60 minutes at room temperature and 4×3 minute washes in TBSMT were conducted between steps. Chromogenic development was performed by BCIP/NBT reaction in Alkaline Phosphatase buffer (100 mM TrisHCl pH 9.5, 5 mM MgSO_4_, 0.01% Tween-20) before quenching the reaction by washing in distilled water followed by TBSMT. For all Coomassie stained gels proteins were separated by SDS-PAGE as above and then stained with SimplyBlue™ SafeStain (Thermo Fisher Scientific) as per manufacturer’s standard protocol.

#### Genotyping

The strains verified above as expressing RBD::mClover were subject to genomic DNA extraction using the Chelex 100 boil method (HwangBo et al., 2010). Primers flanking the transgene were then used to amplify the entirety of the RBD::mClover coding region using Q5 DNA polymerase (New England Biolabs). PCR amplicons were size-verified by electrophoresis on a 0.8% TAE agarose gel stained with SYBR Safe (Thermo Scientific, Waltham, MA) and then excised and purified using the Wizard^®^ SV Gel and PCR Clean-Up System kit (Promega, Madison, WI) according to manufacturer’s protocol. The purified amplicons were then subcloned in to pJet1.2 using the CloneJET PCR Cloning Kit (Thermo Scientific, Waltham, MA) according to manufacturer’s protocol and the ligation product transformed in to DH5alpha chemically competent cells (New England Biolabs, Ipswich, MA) and sequenced using primers FDX1_seq1 and Clov_seq1 (5’-TAGCGCAGCTTCGCCTACAT-3’ and 5’-GCTGAACTTGTGGCCGTTC-3’).

#### Scaled Cultivation and Bulk Protein Purification

Algae cultures were grown in a semi-continuous fashion in 1 L baffled flasks in TAP media wherein 95% of the culture was harvested via centrifuge and replaced with new media every two days; harvested call pellets were snap frozen until lysis. Cells were lysed in 1X Bugbuster (diluted from 10X buffer free stock) buffered with 50 mM Tris-HCl pH8.5 containing 1X Pierce Protease Inhibitor Cocktail (A32955; Thermo Scientific, Waltham, MA), 250 U/mL of Basemuncher endonuclease (ab270049, Abcam, Cambridge, MA), 1 mM DTT, 1 mM EDTA. 10 mL of lysis buffer was added to each gram of snap frozen wet biomass with typically 3-6 grams of wet biomass being processed at once. To the partially clarified cell lysate 10% wt/vol Polyethylenimine (Product#408719, Sigma-Aldrich, St. Louis, MO), adjusted to pH 8.5 by addition of concentrated HCl was added to a final concentration 0.1% wt/vol. The lysate was then centrifuged at 6000 rcf at 4°C for 5 minutes and supernatant recovered. The lysate was then further decolorized by gently shaking against an equal volume of Methyl tert-Butyl Ether twice and then Hexanes (Fisher Scientific). The aqueous layer was then recovered, filtered through a 0.45 μm PES syringe filter and applied to a Capto Q anion exchange resin for column chromatography purification on an Akta pure 150 system fitted with an external sample pump (Cytiva, Amersham, UK).

#### Protein Chromatography

A 5 mL Capto Q resin anion exchange column (Cytiva) was equilibrated with 5 column volumes (CV) of 50 mM TrisHCl pH8.5. The clarified and decolorized lysate was then applied to the column through the system pump at 3 mL/minute. The column was washed with 10 CV equilibration buffer. The RBD::mClover enriched fraction was eluted with 5 CV of 100 mM NaCl in Tris-HCl pH8.5 buffer. During bulk purification ~20 mL of lysate was bound to the column before binding capacity for the RBD was reached. When 40 mL of clarified lysate were processed for bulk purification, 20 mL of lysate was first bound to the column, washed, and eluted as above before the column being immediately stripped by the addition of 5 CV of 1 M NaCl then 5 CV of 1 M NaOH before re-equilibrating with 10 CV 50 mM Tris-HCl and performing a second purification cycle.

A 5 mL Phenyl resin prepacked Hydrophobic Interaction column (Cytiva) was equilibrated with 5 column volumes (CV) of 0.4 M (NH_4_)_2_SO_4_, 50 mM Sodium Phosphate buffer pH 7.4. Immediately after Anion Exchange fractionation, the combined 100 mM NaCl fractions containing the RBD::mClover were brought to 0.4 M (NH_4_)_2_SO_4_ by addition from a 2 M stock solution. The RBD::mClover fraction was then applied to the Phenyl HP resin at a rate of 3 ml/min before being washed with 7.5 CV of 0.35 M (NH_4_)_2_SO_4_, 50 mM Sodium Phosphate buffer pH 7.4. The bound RBD::mClover was then eluted by the addition of 5 CV of 0.15 M (NH_4_)_2_SO_4_, 50 mM Sodium Phosphate buffer pH 7.4. The eluted fraction was then concentrated using Amicon Ultra 15 diaconcentrators with a 10kDa cutoff (Cat # UFC901024, Millipore, Temecula, CA) and buffer exchanged 3 times with PBS pH7,4 (Cat # 10010023 Gibco, Waltham, MA) before concentrating down to 3 mL.

#### ACE2 Receptor Binding assay

Strepavidin coated microtiter plates (#15124, Thermo Fisher) washed 3X in Receptor Assay Blocking buffer (25 mM TrisHCl, 150mM NaCl, pH 7.4, 0.1% wt/vol Bovine Serum Albumin, 0.05% vol/vol Tween-20) and then were coated with 50 ng per well of biotylated human ACE2 produced in HEK cells lines (#10108-H08H-B Sin Biological) dissolved in 100 μL of PBS for one hour at room temperature with gentle shaking on an orbital table. Recombinant SARS-CoV-2 RBD fused C-terminally to rabbit IgG Fc (RBD::rFc, # 40592-V31H Sino Biological) was used as a specific competitor. A dilution series of 100 to 0.01 μg/mL RBD::rFc was made in PBS containing 2 μg/mL of algae produced ER-Golgi retained RBD::mClover including a no-RBD::rFc reference. Wells were washed four times with Receptor Assay Blocking buffer and then filled with the RBD competitor solution at 30 μL/well for 1 hour at room temperature as above. Wells were washed four times and then filled with 100 μL of Goat anti-GFP conjugated to Horseradish Peroxidase (ab6663, Abcam) diluted 1:3000 in Receptor Assay Blocking buffer. After a 30-minute incubation, wells were washed four times and then 100 μL of Pierce TMB substrate kit added. The chromogenic reaction was incubated at room temperature for 20-25 minutes and then quenched with 100 uL of 2 M HCl. The absorption was then measured at 490 nm using a Tecan Infinite m200 Pro plate reader.

#### Characterization by Protein Mass Spectroscopy

Protein lysate from the Chloroplast-directed and ER-Golgi Retained strains were prepared as above and RBD protein purified by Anion exchange chromatography using Capto-Q resin. It should be noted that the ER-Golgi retained version of RBD::mClover eluted at 100 mM NaCl in one fraction when a 100 mM stepped gradient from 0 to 500 mM NaCl was tested, while the Chloroplast directed version eluted over a broad range of salt concentrations from 200 to 400 mM NaCl, suggesting marked differences in affinity to the Capto Q anion exchange resin. The RBD::mClover enriched fractions where then diaconcentrated and buffer exchanged three times against 50 mM Tris-HCl, pH 8.5. These samples were then further purified using Anti-GFP mAb-Magnetic Beads (# D153-11, MBL International, Woburn, MA) according to manufacturer’s protocol except for washing with 50 mM Tris-HCl pH 8.5 with 0.05% Tween-20. Samples were submitted for mass spectrometry at the University of California, San Diego’s Biomolecular and Proteomics Mass Spectrometry Facility as either on-bead digestions or as excised bands from SDS-PAGE gels stained with SimplyBlue Coomassie stain according to manufacturer’s instructions (Thermo Scientific).

## Supporting information

pRMC1,2,3 Vector Maps

