## Supplemental Figure 1 for "Recombinant production of a functional SARS-CoV-2 spike receptor binding domain in the green algae *Chlamydomonas reinhardtii*"

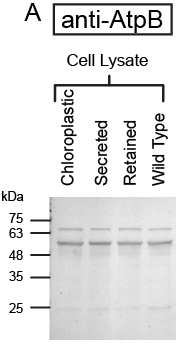

Figure S1. Same amounts of cell lysate protein (5 ug as determined by Bradford assay) were loaded on to an SDS-PAGE gel and Chloroplastic AtpB protein was detected with anit-AtpB antibodies to demonstrate equal loading of solubilized protein. These samples are paired with the cell lysate samples in Figure 2.
